# Plasticity and invariance of Arabidopsis inflorescence and floral shoot apical meristems in response to mineral nutrients

**DOI:** 10.1101/2025.01.31.635844

**Authors:** Benoit Landrein, Katie Abley, Pau Formosa-Jordan, Elliot M. Meyerowitz, Henrik Jönsson, James C. W. Locke

## Abstract

In many species, floral organ production is invariant while flower production rate can be plastic. This allows plants to adapt flower number to their environment whilst maintaining a constant flower structure. The *CLAVATA/WUSCHEL* feedback loop underpins both inflorescence (IM) and floral meristem (FM) activity, respectively responsible for flower and floral organ production. We explore how plasticity and invariance can differ between IM and FM in response to nutrient availability. FM size is less sensitive to changes in nutrients than IM size, and floral organ production is insensitive to these small size changes. However, *clavata3* mutants display larger changes in FM size, approaching those observed in WT IM under nutrient change, with increased floral organ number. This suggests that invariant floral organ production requires that FM size undergoes limited changes in response to nutrients. Compared to the IM, in the FM, levels of cytokinin (CK) signaling are lower and CK signaling and *WUSCHEL* expression are less impacted by nutrient level. Through genetic perturbations, we show a reduced response of FMs to varying cytokinin levels. Our work shows one way that the balance between plasticity and invariance can be set differently in different contexts.

## Introduction

Plants show remarkable levels of developmental plasticity, which can be defined as the same genotype producing a reproducible change in phenotype in response to a difference in the environment (Abley *et al*., 2016). This ability of plants to adapt their shape and physiology to a variety of environmental cues is a key evolutionary adaptation that has allowed these organisms to colonize a large variety of environments (Alpert and Simms, 2002). Molecular sensors are able to perceive environmental signals and specific mechanisms exist to integrate internal and external signals into a developmental output that is appropriate for the given environment (Chaiwanon *et al*., 2016). However, some plant traits are relatively invariant, meaning that the phenotype produced by a genotype is constant and insensitive to differences in the environment (Berg, 1960; Armbruster *et al*., 1999; Abley *et al*., 2016). For a given species, dimensions of floral organs often remain invariant even when other traits such as leaf size, leaf number and stem height vary in a correlated way between conditions (Berg, 1960; Armbruster *et al*., 1999). This raises the question of how developmental mechanisms differ between plastic and invariant traits.

In a previous study of the mechanisms underlying developmental plasticity, we showed that high nitrate level in the soil increases the rate of flower production by increasing the number of stem cells in structures called shoot apical meristems (Landrein *et al*., 2018). Although flower production rate varies according to the environment, flower structure tends to be invariant. Within a *Brassicaceae* species such as Arabidopsis, the number of floral organs per whorl is usually fixed irrespective of the environment (Nikolov, 2019). Here we compare the mechanisms that control the rate of flower production (a plastic trait) and the number of floral organs (an invariant trait) and their respective sensitivities to changes in nutrient levels. To do this, we compare the two types of shoot meristem involved in flower and floral organ production: inflorescence meristems (IM), and floral meristems (FM). Inflorescence meristems are indeterminate structures found at growing shoot apices and themselves produce FMs from their flanks (Barton, 2010). IMs exhibit spiral phyllotaxis and each new FM is generated periodically and laid down at approximately the golden angle (137.5°) with respect to the previous one (Landrein *et al*., 2015). In contrast, the Arabidopsis FM has whorled phyllotaxis, in which all organs of each type (sepals, petals, stamens, carpels) are produced nearly simultaneously in a ring around the meristem (Smyth *et al*., 1990).. The primary IM produces new flower primordia continuously for approximately two weeks, whilst the FM is determinate and produces four whorls of organs before its development terminates (Smyth *et al*., 1990; Bowman *et al*., 1991; Lenhard *et al*., 2001; Lohmann *et al*., 2001)

Meristems contain a pool of post-embryonic stem cells whose activity allows organogenesis to occur during the entire life of the plant. In both IMs and FMs, stem cells are located in the central zone, a small region of cells at the apex of the meristem (Meyerowitz, 1997; Barton, 2010). Stem cell fate and differentiation are regulated through the activity of a negative feedback loop involving *WUSCHEL* (*WUS)* and *CLAVATA* (*CLV*) genes. *WUS* encodes a homeodomain transcription factor expressed below the stem cell niche, in the organizing center, that can move to the central zone through plasmodesmata (Laux *et al*., 1996; Mayer *et al*., 1998; Yadav *et al*., 2011; Daum *et al*., 2014). In the central zone, WUS directly induces the expression of *CLAVATA3* (*CLV3*), which codes for a mobile extracellular peptide that diffuses to the organizing center to repress *WUS* expression, through binding to specific receptors, including notably *CLV1* (Clark *et al*., 1993; Clark *et al*., 1995; Fletcher *et al*., 1999; Schoof *et al*., 2000; Ohyama *et al*., 2009). *WUS* also represses the expression of *CLV1* (Busch et al., 2010), and CLV3 binding causes internalization and degradation of CLV1 (Nimchuk *et al*., 2015), thus adding additional feedbacks to this network. This negative feedback between *WUS* and the C*LV* complex is thought to be central for the maintenance of the stem cell niche in the shoot apical meristem, with the negative regulation of *WUS* by CLV3 being key to controlling meristem size (Schoof *et al*., 2000; Brand *et al*., 2000).

Cytokinin (CK) hormones are involved in the regulation of stem cell homeostasis in the SAM as they can induce *WUS* expression and may also position its expression domain in the organizing center (Gordon *et al*., 2009; Chickarmane *et al*., 2012; Gruel *et al*., 2016). CK also directly affects cell division in the SAM, notably through MYB3R4 (Yang *et al*., 2021). In return, WUS also induces cytokinin signaling through the repression of negative regulators from the type-A ARR (ARABIDOPSIS RESPONSE REGULATORS) family thus generating a positive feedback loop between WUS and cytokinins (Leibfried *et al*., 2005; Gordon *et al*., 2009; Busch *et al*., 2010). Cytokinins are involved in the SAM response to the environment and cytokinin levels in the plant provide a systemic readout of nitrate availability (Landrein *et al*., 2018; Asim *et al*., 2020; Abualia *et al*., 2023). In the vegetative meristem of germinating seedlings (which produces leaves), light and metabolic signals have been shown to influence *WUS* expression through CK degradation (Pfeiffer *et al*., 2016). In the IM, nitrate can induce *WUS* expression and thus affect stem cell homeostasis through the generation of a long-range signal of cytokinin precursors (Landrein *et al*., 2018). As increasing stem cell number also leads to an increase in meristem size and organogenesis rate (Schoof *et al*., 2000; Landrein *et al*., 2015), which may result from WUS regulation of auxin signaling (Ma *et al*., 2019), the CK-dependent induction of *WUS* expression in the inflorescence meristem by nitrate can act as a mechanism to control flower production rate in response to nitrate availability (Landrein *et al*., 2018).

These recent studies have emphasized the central role of the core stem cell regulator *WUS* in the response to environmental signals of two types of SAM, the vegetative and the inflorescence meristem. As floral meristems also contain pools of stem cells whose activity is controlled by the core *WUS/CLV* feedback loop (Clark *et al*., 1993, 1995; Laux *et al*., 1996), a key question is how the invariance of floral development can be maintained in the face of environmental input into *WUS*. To study the differences in invariance and plasticity in response to environmental inputs into organ production in different types of meristems, we performed a comparative analysis of the response of floral and inflorescence meristems to mineral nutrients and to mutations that affect meristem size.

We show that floral meristem size and organ number are less sensitive to nutrient availability than inflorescence meristem size and flower production rate. The robustness in floral organ number is partly due to the floral meristem’s resilience to small size changes, requiring larger increases to affect organ number compared to the inflorescence meristem. In *clv3* mutants, where floral meristem size increases similarly to the inflorescence meristem under nutrient-rich conditions, this invariance is lost, leading to changes in organ number. In wild-type plants, large increases in floral meristem size are avoided because it is less sensitive to nutrients than the inflorescence meristem, contributing to the invariance of floral organ number. We find that the reduced sensitivity to nutrients in the floral meristem may be related to its six-fold lower levels of CK signaling. Floral meristem size and organ number are also almost unaffected by genetic disruptions in the cytokinin pathway, which we previously found to be essential for the increase in WUS expression and meristem size in the inflorescence meristem when mineral nutrients are added (Landrein et al., 2018). Thus, floral organ number invariance relies on nutrient insensitivity in the floral meristem.

## Results

### Floral and inflorescence meristems respond differently to changes in mineral nutrition

To study the effect of mineral nutrition on the robustness and plasticity of organogenesis in Arabidopsis inflorescence meristems (IMs) and floral meristems (FMs), we analysed wild-type (WT) plants grown on soil or on a mix of soil and sand to reduce mineral nutrient availability (Landrein *et al*., 2018). Although these different growing substrates could have effects on plant growth through their different densities and water contents, we previously showed that the two conditions caused similar changes to IM size and organogenesis to those seen under different nitrate concentrations (Landrein *et al*., 2018). We measured the number of flowers produced by the primary inflorescence meristem within 10 days after bolting and the number of floral organs produced by FMs. As already characterised (Landrein *et al*., 2018), plants grown on soil produced more flowers during the development of the primary inflorescence than plants grown on a mix of soil and sand (Fig. 1A and B). The coefficients of variation of flower number produced within 10 days of bolting (standard deviation divided by mean), which provide a measure of the variability in the trait, were 0.13 for the mixture of soil and sand and 0.12 for soil. We then looked at the effect of the nutritive conditions on floral organ production by the FMs. In agreement with the idea that floral organ number is fixed in many *Brassicaceae* (Nikolov, 2019), we observed almost no variability within each condition and no effect of the growth conditions on the number of floral organs in WT Arabidopsis plants. Most of the flowers showed the classical pattern of 4 sepals, 4 petals, 6 stamens and 2 carpels (Fig. 1A and C). The coefficient of variation of the number of stamens, that occupy the most variable floral whorl, was low in both conditions: around 0.06 in the mixture of soil and sand and around 0.07 in soil. These quantifications show that floral organ number is invariant within conditions and not plastic between conditions, while flower number produced within 10 days of bolting is more variable within conditions and plastic between conditions.

**Fig. 1.**
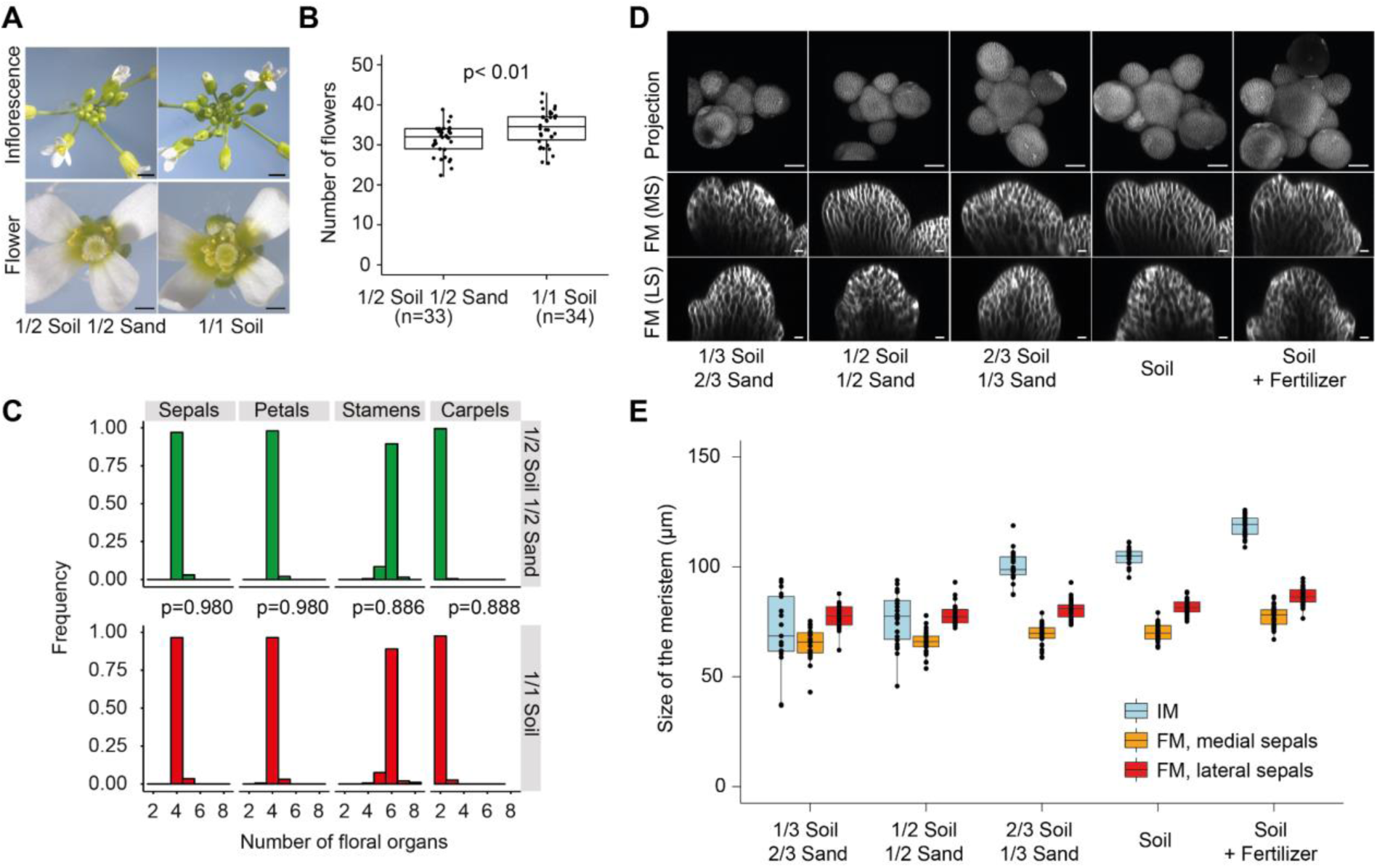
Nutrients affect inflorescence and floral meristems differently in WT plants. **A.** Representative inflorescences (top) and flowers (bottom) produced by Col-0 WT plants growing on a mix of soil and sand or on soil only. Scale bars, top: 1 cm, bottom: 0.1 cm **B.** Number of flowers produced by the primary inflorescence within the first 10 days of flowering by plants growing on a mix of soil and sand or on soil only. Pool of 2 independent experiments. The effect of the growth conditions was assessed using a bilateral Student test, revealing a significant difference (p<0.01). **C.** Number of floral organs produced by plants growing on a mix of soil and sand or on soil only. 200 flowers from 20 plants from 2 independent experiments were sampled for each condition. The effect of the growth conditions was assessed using linear mixed-effects models. **D.** Stack projections of representative IMs (top) and sections through the middle of the medial (middle) and lateral sepals (bottom) of representative FMs produced by plants growing on soils of different nutritive qualities. Scale bars, top: 50 µm, middle and bottom: 10 µm. **E.** Size of IMs and stage 3 FMs (measured as the distance between the medial sepals or between the lateral sepals, see Supplementary Fig. S1) produced by plants growing on soils of different nutritive qualities. 17 to 26 IMs and 23 to 43 FMs from 2 independent experiments were used for each condition.The data on flower number (B) and IM size (D and E) were previously generated and analysed in (Landrein *et al*., 2018). The data on FM size in E was previously generated, but not analysed, in (Landrein *et al*., 2018). The top panel of D is equivalent to part of Fig. 1A of (Landrein *et al*., 2018) and the IM data in E is the same as that plotted in (Landrein *et al*., 2018), Fig. 1C.

The increase in flower production rate we observed in a higher nutrition environment has been previously linked to an enlargement of the inflorescence meristem (Landrein *et al*., 2018). We thus compared the effect of mineral nutrients on FM and IM diameters by reanalyzing a previous set of data where we imaged meristems produced by plants growing on 5 different types of soil (Landrein *et al*., 2018). We measured the diameter of the IM using a previously developed method (Landrein *et al*., 2018) and the diameter of the floral meristems by measuring the distance between the medial (abaxial and adaxial) and between the lateral sepals on confocal sections of stage 3 flowers (Smyth *et al*., 1990; McKim *et al*., 2017) (Supplementary Fig. S1A and B). We observed a positive correlation between the diameter of the IM and the availability of mineral nutrients in the soil (Fig. 1D-E) (linear models fitted to the data revealed the effect of the nutrient condition on IM diameter was significant in both experimental replicates (p < 0.001 in both cases)). Accordingly, we measured that plants growing in the highest nutritional condition showed, on average, a 68% increase in the diameter of the IM compared to plants growing in the lowest nutritive condition (58% increase in replicate 1, 77% increase in replicate 2). Conversely, we observed that mineral nutrients had less effect on the diameter of the floral meristems as plants growing in the highest nutritional condition only showed a 20% increase in the diameter of the FM when measured as the distance between medial sepals (20% for replicate 1, 19 % for replicate 2) and a 12% increase when measured as the distance between the lateral sepals compared to plants growing in the lowest nutritive condition (16% increase for replicate 1, 9% for replicate 2) (Fig. 1D-E, Supplementary Fig. S1D, F). This smaller effect of nutrient condition on floral meristem diameter was however statistically significant (linear model, p < 0.001 for the distance between the medial sepals and for the distance between the lateral sepals in both experimental replicates). Because flower production is thought to depend on auxin transport in the epidermis (Jönsson *et al*., 2006), we also measured the distance between medial and lateral sepals following the epidermis (Supplementary Fig. S1B) and observed a similar effect of the growth conditions on floral meristem size (Supplementary Fig. S1C). The diameters of the IMs and FMs were correlated across plants and nutrient conditions (Supplementary Fig. S1D) and the diameters of both meristem types were correlated with shoot weight (Supplementary Fig. S1E). From these experiments, we conclude that both the IM diameter and the FM diameter are positively correlated with nutrient availability, which could indicate similar regulatory mechanisms operating in both contexts, but floral meristem diameter and floral organ number are less affected by nutrition than inflorescence meristem diameter and number of flowers.

### Floral organ number correlates with meristem size in *clv3* mutants

We next tested whether floral organ number is sensitive to larger changes in meristem size, or if it is invariant to any change in meristem size. It is indeed possible that the different phyllotactic pattern in the FM, or other differences in regulation between the FM and IM, causes the FM organ number to be invariant to meristem size. To test this, we looked at the relationship between meristem diameter and organ production in IMs and FMs of *clv3* mutants, which have larger floral meristems than wild-type plants grown in nutrient rich conditions. The negative feedback loop between *CLV3* and *WUS* is known to restrict *WUS* expression and allow tight control of stem cell proliferation during development (Schoof *et al*., 2000). Accordingly, inhibiting CLV expression leads to an over-proliferation of the stem cells, which affects meristem size and organ production in both inflorescence and floral meristems (Clark *et al*., 1993, 1995). We investigated the relationship between FM diameter and the number of floral organs produced under different nutritive conditions in two mutant alleles of *clv3* in the Col-0 ecotype: *clv3-7* and *clv3-9* (Fletcher *et al*., 1999; Nimchuk *et al*., 2015).

In both growth conditions (½ soil ½ sand and soil), mutations in *CLV3* led to a strong enlargement of the inflorescence meristem, often leading to fasciation (which precludes precise measurements of IM size in *clv3* mutants), especially in *clv3-9*, whose mutant phenotype was stronger than that of *clv3-7* (Fig. 2A). In the flower, we observed that mutations of *CLV3* also led to a strong enlargement of the floral meristems with *clv3* FMs having significantly larger diameters than WT FMs in both nutrient conditions (*clv3-7* is 20% larger than WT in soil and 17% larger in ½ soil ½ sand; *clv3-9* is 27% larger than WT in soil and in ½ soil ½ sand, all p values < 0.001 from the linear model t-tests, Fig. 2C and Supplementary Fig. S2A). On average, across the experimental replicates, we observed a similar response of FM diameter to mineral nutrients in *clv3* and in the WT (Fig. 2C) (9.8% increase for *clv3-7*; 7.4% increase for *clv3-9*; and 7.7% increase in the WT; p value for the interaction between genotype and nutrient in the linear model t-tests was non significant (p>0.05) for both alleles).

**Fig. 2.**
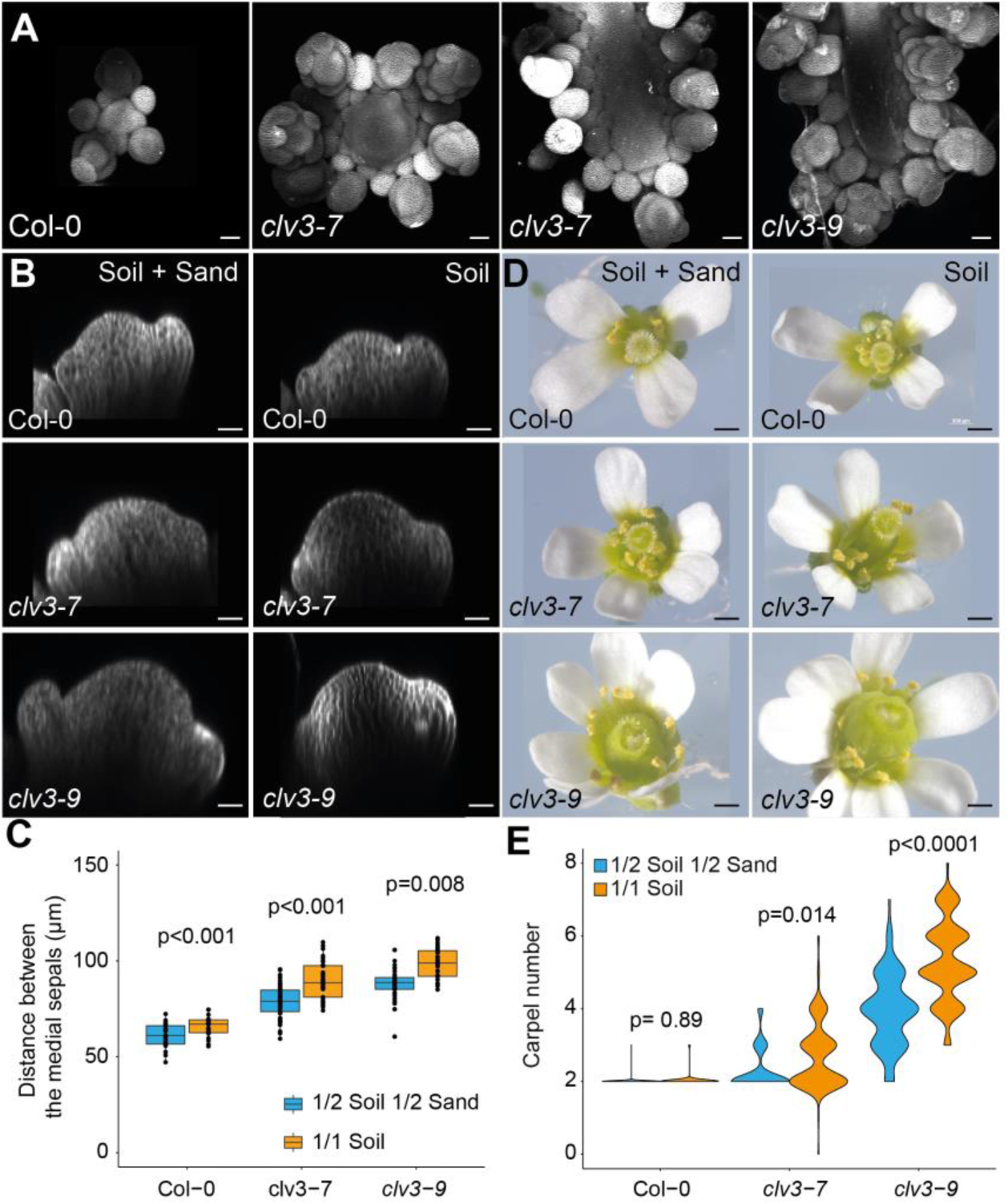
Nutrition affects floral meristem size and floral organ number in *clv3* mutants. **A.** Representative inflorescence meristems produced by Col-0 WT and *clv3* mutant plants growing on soil. Two examples of *clv3-7* inflorescence meristems with different sizes are shown. Note that the right-most pictures of *clv3*-*7* and *clv3*-*9* only show part of a fasciated meristem. Scale bars, 50 µm. **B.** Section through the middle of the medial sepals of representative stage 3 floral meristems in Col-0 and *clv3* mutants from plants grown on a mix of soil and sand or on soil only. **C.** Distance between the medial sepals of stage 3 flowers produced by Col-0, *clv3-7* and *clv3-9* grown in different nutrient conditions. 11 to 24 plants from 2 independent experiments were sampled for each genotype and condition. A linear mixed-effect model was used to assess the effect of the growth conditions. **D.** Representative flowers of Col-0 and *clv3* flowers produced by plants grown on a mix of soil and sand or on soil only. Scale bars, 0.5 mm. **E.** Violin plot representation of the frequency of carpels produced by Col-0 and *clv3* mutant flowers growing on soil. 200 flowers from 20 plants from 2 independent experiments were sampled for each condition. A linear mixed-effect model was used to assess the effect of the growth conditions.

We also characterised the relationship between meristem diameter and floral organ number for the FM and compared this with the relationship between IM diameter and the rate of organ production for the IM in WT and in *clv3* mutants. We found that changes in meristem diameter between the 50:50 mixture of soil and sand and the soil condition were associated with changes in flower number produced within a 10-day period in the IM but not with changes in floral organ number in the FM (Fig. 1). To investigate this further, we looked at the correlation between organogenesis rate and meristem diameter in the IM by measuring the plastochron ratio (i.e. the mean difference in size between successive primordia ordered following the spiral pattern of phyllotaxis for each IM (Landrein *et al*., 2015)). Measurements of the plastochron ratio are based on the assumption that the difference in size between successive organs should be proportional to the time delay between successive initiations, and should thus be inversely proportional to the organogenesis rate. As we previously observed while measuring IMs of different sizes (Landrein *et al*., 2015), the plastochron ratio strongly correlated with the diameter of the IM in WT plants grown on soils of different nutritional qualities but also in plants showing alteration in CK metabolism (Fig. 3A and Supplementary Fig. S3A). In the IM, organogenesis rate is thus directly correlated with meristem diameter so that even small changes in meristem diameter lead to changes in flower production rate.

**Fig. 3.**
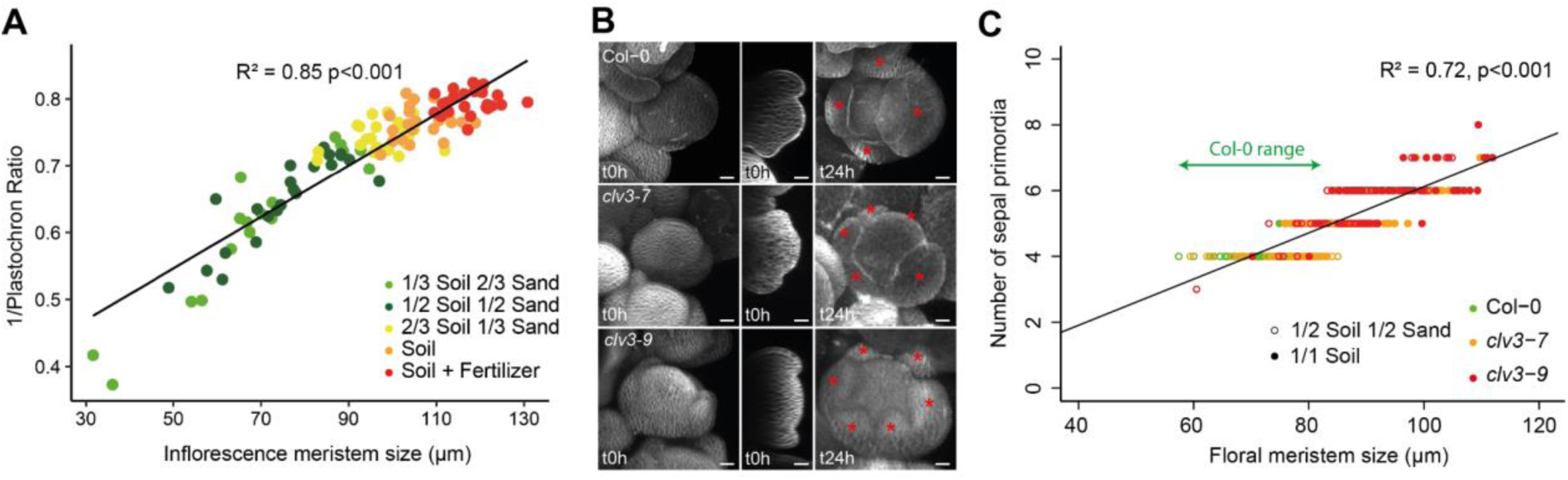
Organ number correlates to meristem size in both inflorescence and floral meristems. **A.** Correlation between the organogenesis rate (measured as the inverse of the plastochron ratio) and the size of the inflorescence meristem in Col-0 WT plants growing in soil of different nutritive qualities. 17 to 26 IMs from two independent experiments were sampled per condition. A linear model was applied to fit the data. **B.** Live imaging of floral meristems obtained through *in vitro* SAM culture and showing the emergence of supernumerary sepals in *clv3* mutants. Sepals at 24h are marked with asterisks. Scale bars, 20µm. **C.** Correlation between the number of sepal primordia and the size of the floral meristems (obtained by measuring the distance between the medial sepals) in a pool of Col-0 and *clv3* mutant plants growing on a mix of soil and sand or on sand only and cultivated *in vitro* for 24h. 11 to 24 plants from 2 independent experiments were sampled for each condition. A linear model was applied to fit the data. Col-0 floral meristem size range is overlaid in green. The data on IM size and plastochron ratio (A) were previously generated and analysed in (Landrein *et al*., 2018).

In the FM, we know that this correlation does not hold because the small changes in meristem diameter that are induced by modulations of the growth conditions do not lead to changes in floral organ number (Fig. 1). However, larger increases in meristem diameter, obtained when *CLV3* expression is affected, led to the production of more floral organs (Fig. 2 and Supplementary Fig. S2B-D). Floral organ number was affected by the growth conditions in *clv3* so that the number of floral organs was increased when plants were grown in nutrient rich conditions (Fig. 2E, Supplementary Fig. S2B, D). This increased response of floral organ number to nutrients in *clv3* compared to Col-0 was significant for the number of carpels produced in *clv3-9* (p value for the interaction between genotype and environment in a Poisson regression model <0.001). We also observed that floral organ number was more variable within conditions in *clv3* mutants than in the WT (the coefficients of variation of floral organ number being between 0.13 and 0.33 for *clv3* depending on the organ type and the growth conditions but only up to 0.08 in the WT, Fig. 2D and Supplementary Fig. S2B-D). We could not detect any effect of the *clv3* mutation on FM diameter variability (the coefficient of variation being close to 0.1 in both WT and *clv3* mutants). Overall, these results suggest that *CLV3* activity is necessary for the invariance of floral organ numbers within and between conditions in Arabidopsis.

We next tested whether floral organ number correlated with floral meristem diameter in *clv3* mutants. To do so, we performed time-lapse microscopy on developing flower buds following published protocols (Fernandez *et al*., 2010) so that we could measure the diameter of the floral meristems at stage 3, let the primordia develop, and measure accurately the number of sepal primordia at stage 4, when all of the sepals are fully emerged (Fig. 3B). Pooling all WT and *clv3* data together, we observed a strong correlation between the number of sepal primordia and the diameter of the floral meristems, assessed through the distance between the medial sepals (Fig. 3C and Supplementary Fig. S3B). We also observed that floral meristems needed to reach a certain diameter for more than 4 sepal primordia to emerge. The mean FM diameter in Col-0 across both nutrient conditions was approximately 72 µm (as measured between the medial sepals). Although the distributions of *clv3* mutant FM diameters for different sepal numbers overlap, when FM diameters increased above 85 µm in *clv3* mutants, only flowers with more than 4 sepals were observed (Fig. 3C). The correlation between the number of sepal primordia and FM diameter did not hold true when meristem diameter was assessed through the distance between the lateral sepals, which is probably due to the fact that floral meristems fasciate along the medial axis in *clv3* mutants, thus losing their radial symmetry (Supplementary Fig. S3C and D).

Thus, although the FM is robust to small changes in meristem size in col-0, larger meristem sizes allow a new organ to be inserted. This is in contrast to the IM, where small increases in meristem diameter lead to increases in the rate of organ production. However, this difference between the two types of meristem does not alone explain why the number of floral organs remains invariant within an environment and under different nutritive conditions. Indeed, we observed that when the floral meristem diameter increased above 85 *µ*m in *clv3* mutants there was a consistent increase in floral organ number (Fig. 3C). This represents an approximately 23% increase in floral meristem diameter from a mean of around 69 *µ*m in WT on the 50:50 mix of soil and sand. We previously observed only a 20% increase in WT FM diameter across the full range of nutrients (from ⅓ soil ⅔ sand to soil + fertilizer) with WT plants rarely exceeding a floral meristem diameter (measured between the medial sepals) of 85*µ*m (Fig. 1, S1D, E, 3C). If floral meristem diameter in WT had responded to mineral nutrients as much as the inflorescence meristem, which showed around 37% increase in diameter between the 50:50 mixture of soil and sand and soil (Fig. 1), compared to only a 7.7% increase in FM diameter in this experiment (and a 6.5 % increase in the experiment shown in Fig. 1), this would have resulted in the FM diameter reaching around 95 *μ*m on soil. According to the relationship between floral meristem diameter and floral organ number obtained by including both Col-0 and *clv3* mutants, a FM diameter of around 95*μ*m would be predicted to result in a shift in the number of sepals from 4 to 5 or 6 (Fig. 3C). These findings suggest that although the FM requires larger increases in meristem size for an increase in organ number, the decreased response of the FM diameter to nutrients also plays a role in the invariance of floral organ number. We thus next investigated the mechanisms underlying this reduced sensitivity of the FM to nutrients.

### Cytokinin response and *WUS* expression are less affected by mineral nutrition in FM than in IM

Cytokinin hormones play a central role in the regulation of inflorescence meristem function. This is because mineral nutrients trigger the production of cytokinin precursors that move through the vasculature towards the meristem where they are converted to active hormone, leading to increased *WUS* expression and an enlargement of the *WUS* domain of expression, as well as an increase in the size of the IM (Osugi *et al*., 2017; Landrein *et al*., 2018). We thus looked to see if the differences in meristem response to nutrients that we observed between inflorescence and floral meristems could be due to differences in cytokinin response and *WUS* expression. We first compared the expression levels (total and maximum signal) and size of the domain of a reporter of cytokinin response (*pTCSn::GFP*) and of a transcriptional reporter for *WUS* (*pWUS::GFP*) in plants grown under different nutritional conditions through a semi-automated pipeline we developed to measure gene expression in both inflorescence and floral meristems (Material and Methods).

For cytokinin response, we observed that both the intensity of the signal and the size of the expression domain of *pTCSn::GFP* were reduced in the FM compared to the IM (Fig. 4A-B, Supplementary Fig. S4A, C). The total signal of *pTCSn::GFP* was approximately 6-fold lower for the FM compared with the IM (Fig. 4B, Supplementary Fig. S4Ci), although the signals of the FM and the IM were correlated across plants and nutritive conditions (Supplementary Fig. S4C i). Despite the reduced expression of *pTCSn::GFP* in the FM, both the total *pTCSn::GFP* signal and the domain size were still responsive to changes in nutrients, although to a lesser extent than in IM (Fig. 4A-B, Supplementary Fig. S4A,C, Supplementary Fig. S5A). Accordingly and depending on the replicate, we measured a 195 to 198 % increase in the total signal and a 31 to 37 % increase in the size of the expression domain of *pTCSn::GFP* between the lowest and highest nutritive conditions in IM and an 81 to 155 % increase in the total signal and only a 9 to 12% increase in the size of the expression domain in FM (Supplementary Fig. S5Ai, ii). For *WUSCHEL*, we observed a smaller difference in signal and in expression domain size between IM and FM, especially in the lowest-nutritive conditions (Fig. 4C and D, Supplementary Fig. S4B). We also observed that mineral nutrition affected *WUS* expression almost similarly in IM and in FM as we measured a 76 to 83% increase in IM and 42 to 68% increase in FM of the signal of the *pWUS::GFP* between the lowest and the highest nutritive conditions (Supplementary Fig. S5Bi). There was also a positive correlation between *pWUS::GFP* in the FM and the IM across all the plants and the nutritive conditions (Supplementary Fig. S4Di). However, we observed stronger differences when looking at the size of the expression domain as we observed a 26 to 35% increase in IM but only a 4-6% increase in the expression domain size of *pWUS::GFP* in the FM between the lowest and the highest nutritive conditions (Supplementary Fig. S5Bii). Also, the *pWUS::GFP* expression domain size was only weakly correlated between IMs and FMs (Supplementary Fig. S4D ii).

**Fig. 4.**
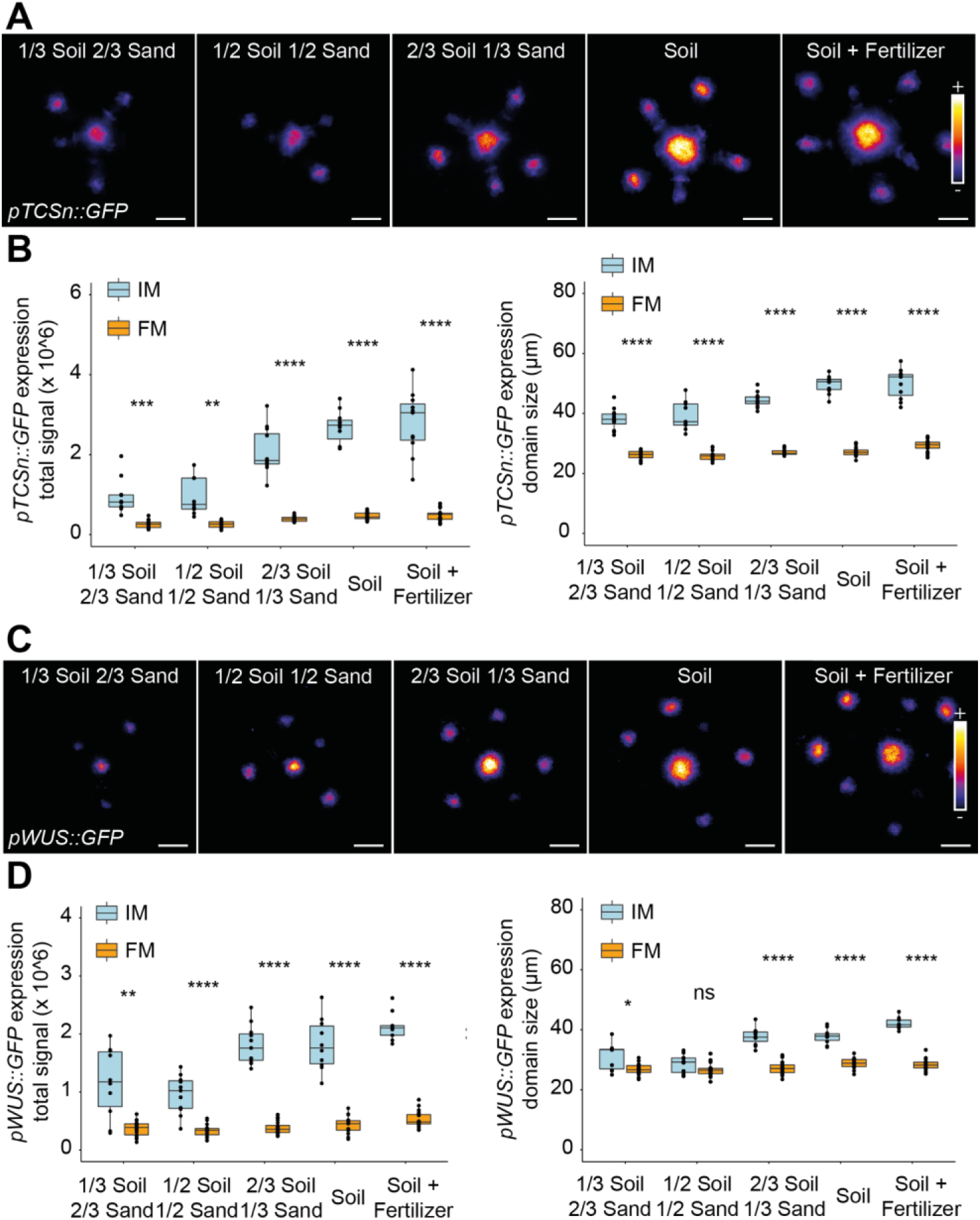
Nutrients affect CK response and WUS expression in IM and FM differently in WT plants. **A.** Representative IMs and FMs of Col-0 WT plants expressing *pTCSn::GFP* (color-coded intensity) produced by plants growing on soils of different nutritive qualities. Scale bars, 50 µm. **B.** Total signal (left) and characteristic diameter of the expression domain (right) of the *pTCSn::GFP* marker in stage-3 FMs and in IMs obtained from plants growing on soils of different nutritive qualities. 13 to 22 FMs produced by 9 to 12 IMs from one experiment were used for each condition (see Supplementary Fig. S4A for another independent experiment). **C.** Representative IMs and FMs of Col-0 WT plants expressing *pWUS::GFP* (color-coded intensity) produced by plants growing on soils of different nutritive qualities. Scale bars, 50 µm. **D.** Total signal (left) and characteristic diameter of the expression domain (right) of the *pWUS::GFP* marker in stage-3 FMs and in IMs obtained from plants growing on soils of different nutritive qualities. 14 to 24 FMs produced by 9 to 11 IMs from one experiment were used for each condition (see Fig. S4B for another independent experiment). Meristem types were compared using bilateral Student tests. Total fluorescence signal measurements (in B and D) are in arbitrary units. The data in A-D were all previously generated and analysed for the IM but not for the FM in (Landrein *et al*., 2018).

These observations suggest that mineral nutrition still affects cytokinin response and WUSCHEL expression in FM but to a lesser extent than in IM, and with lower overall levels of cytokinin signaling in the FM than the IM. The size of the expression domain of *WUS* in FM, which was shown to correlate with meristem diameter in inflorescences (Gruel *et al*., 2016) seems to be relatively insensitive to changes in mineral nutrition.

### Meristem diameter and activity is less affected by cytokinins in the FM than in the IM

Our observations suggest that cytokinins may play a lesser role in setting meristem diameter in the FM than in the IM and thus convey differently the response to mineral nutrients in both types of meristems. To test this hypothesis, we looked at the influence of CK on IM and FM function by characterizing the meristematic defects of mutants affected in CK metabolism. Increasing CK levels in the *ckx3.5 (ckx3; ckx5)* mutant, whose associated genes normally code for members of the CYTOKININ OXIDASE (CKX) family of CK degrading enzymes, caused a 24% increase in IM diameter but only a 5% increase in FM diameter when measuring the distance between the medial or the lateral sepals (Fig. 5A-B and Supplementary Fig. S6A). Additionally, decreasing CK levels in the *ipt3.5.7 (ipt3; ipt5; ipt7*), *log4.7 (log4; log7*) and *log1.3.4.7 (log1; log3; log4; log7*) mutants, whose associated genes normally code for members of the ISOPENTENYL TRANSFERASE (IPT) and LONELY GUY family of cytokinin biosynthetic enzymes, respectively caused 8%, 26% and 42% decreases in IM diameter, and 11%, 13% and 16% decreases in FM diameter when measuring the distance between medial sepals and a 7%, 5% and 12% decrease in FM diameter when measuring the distance between the lateral sepals (Fig. 5A-B and Supplementary Fig. S6A).

**Fig. 5.**
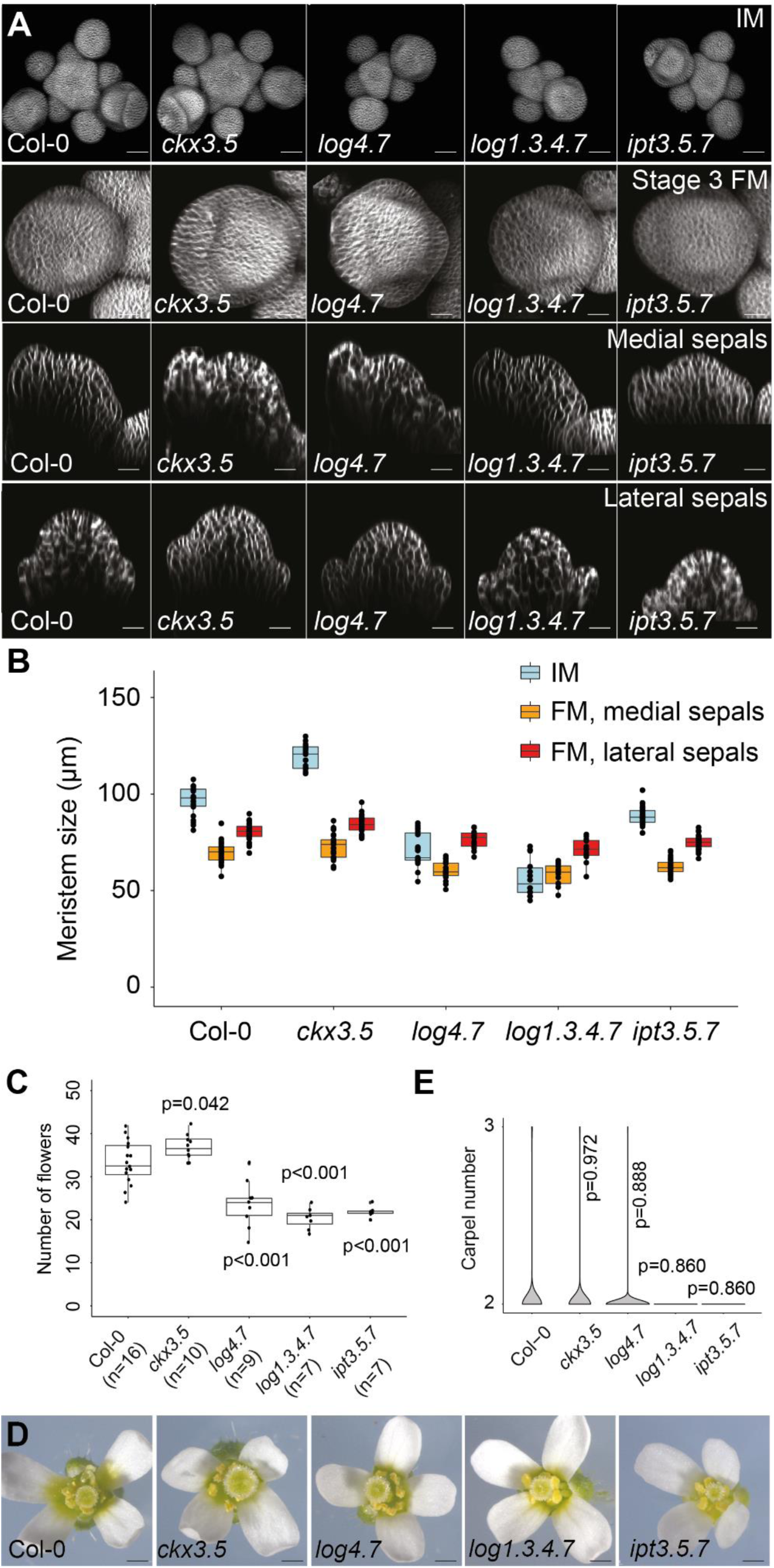
Altering CK metabolism affects inflorescence and floral meristems differently. **A.** Representative IMs and stage 3 FMs (top: projection, middle: section through the middle of the medial sepals and bottom: section through the middle of the lateral sepals) produced by Col-0 WT and mutant plants altered in CK metabolism and growing on soil. Scale bars, 50 µm for the IMs and 10 µm for the FMs. **B.** Size of IMs and stage 3 FMs (measured as the distance between the medial sepals or between the lateral sepals) produced by Col-0 WT and mutant plants altered in CK metabolism and growing on soil. 12 to 41 FMs produced by 14 to 26 IMs from 2 independent experiments were used for each condition. **C.** Number of flowers produced within the first 10 days of flowering by Col-0 WT and mutant plants altered in CK metabolism and growing on soil. One experiment. Each mutant was compared to the WT using a bilateral Student test. **D.** Representative flowers produced by Col-0 WT and mutant plants altered in CK metabolism and growing on soil. Scale bars, 0.1 cm. **E.** Violin plot representation of the number of carpels produced by mutants altered in CK metabolism and growing on soil. 200 flowers from 20 plants from 2 independent experiments were sampled for each condition. Each mutant was compared to the WT using a linear mixed-effect model. The data in B were previously generated and analysed for the IM but not for the FM in (Landrein *et al*., 2018).

These observations thus suggest that FM size may be less affected by changes in CK metabolism than IM size although IM and FM diameters were reduced by a similar small amount in the *ipt3-5-7* mutant. This could be due to the fact that *IPT*s, which are mainly expressed in the vasculature of the root, are generating a systemic signal of CK precursors, while LOGs and CKXs are thought to be mainly expressed in both FMs and IMs where they affect CK metabolism more locally (Bartrina *et al*., 2011; Gruel *et al*., 2016; Osugi *et al*., 2017; Landrein *et al*., 2018). As previously reported, the changes in IM size affect flower production rate in these mutants (Fig. 5C) (Landrein *et al*., 2018). However, the mutants affected in cytokinin metabolism did not show any statistically significant alteration in the number of floral organs in our growth conditions (Fig. 5D,E and Supplementary Fig. S6B) although a small and statistically significant increase in floral organ number has been reported in *ckx3.5* mutant previously (Bartrina *et al*., 2011). Taken together, these experiments suggest that the lack of response of FM size and floral organ numbers to mineral nutrition could be due to the FM’s relative insensitivity to cytokinins.

## Discussion

We compared the response of inflorescence (IM) and floral (FM) meristems to mineral nutrients. We observed that FM diameter was much less affected by the growth conditions than the IM diameter. Accordingly, floral organ production was not affected by mineral nutrient availability, whilst flower production rate was affected. Our work sheds light on processes contributing to the relative invariance of floral organ number: differing relationships between meristem diameter and organ number in the FM and IM; the insensitivity of the FM diameter to cytokinin and mineral nutrients, which is associated with much lower levels of CK signaling in the FM; and the CLV3-mediated negative feedback constraining FM size.

Firstly, we show that FMs and IMs have different relationships between meristem diameter and organ production. The number of organs produced responds similarly in inflorescences and flowers to large changes in meristem diameter but differently to small changes. Small changes in FM diameter do not affect floral organ number while larger alterations, as observed in *clv3* mutants, can lead to changes in the number of organs per whorl. In contrast, the IM shows a continuous relationship between meristem diameter and organ production rate and thus is affected by both small and large changes (Fig. 3A, 6A). These differences may be related to the different types of phyllotaxis in the two types of meristem. The FM only produces a few floral organs in a whorled pattern. To add organs to a whorled pattern would require substantial added space which may be available in the larger *clv3* FMs. In contrast, the spiral phyllotaxis of the IM produces a large number of flowers that appear sequentially and small increases in meristem diameter lead to an increased rate of organ production (Douady and Couder, 1996). Indeed, in contrast to the invariant floral organ numbers in whorled flowers of Arabidopsis, there is often variation in the total organ number in species where spiral phyllotaxis occurs in flowers (Soltis *et al*., 2009; Wang *et al*., 2015), which can correlate with floral meristem size (Wang *et al*., 2015; Kitazawa, 2021). More work will be needed to investigate the hypothesis that the type of phyllotaxis contributes to the level of invariance *versus* plasticity of organ production and to better understand the mechanisms underlying the different relationships between meristem diameter and organ number in whorled *versus* spiral phyllotactic patterns. One possibility would be to test whether species with spiral phyllotaxis in flowers have increased plasticity in floral organ number in response to nutrient conditions.

However, the differences in the relationship between meristem diameter and floral organ number between the FM and the IM cannot alone account for the contrast between the invariance of floral organ number and the plasticity of flower production rate under varying nutritive conditions. FM diameter is much less responsive to nutritive conditions than IM diameter, and our *clv3* results reveal that when the FM diameter increases (due to this genetic perturbation) to an extent that approaches the increase in IM diameter in response to nutrients, extra floral organs are produced (Fig. 3). Thus, the reduced response of FM size to nutrients is likely important for understanding the invariance of floral organ number.

We show that while the IM responds to changes in CK conditions and to mineral nutrients, the lack of response of the FM to mineral nutrients correlates with a relative insensitivity to cytokinins, as assessed by the impact of mutations that affect CK levels. There are multiple possible explanations for the reduced sensitivity of the FM to perturbations in CK levels. One could be that the differences between the FM and IM in the CK response are due to differences in the expression or regulation of genes involved in CK metabolism. For instance, while *LOG4* mediates the production of active CK in the L1 of the IM, it is LOG7 that mediates the production of CK in the FM (Gordon *et al*., 2009; Chickarmane *et al*., 2012; Gruel *et al*., 2016)). However, the expression of the *pTCSn::GFP* reporter in FMs was correlated with that in IMs across plants and nutritive conditions (Supplementary Fig. S4C), perhaps indicating similar mechanisms of regulation of CK response in both tissue contexts, although with a reduced magnitude of the response in the FM. Another possibility is that, rather than being solely related to reduced responsiveness of CK signaling to nutrients, the reduced sensitivity of FM size to CK is related to reduced levels of CK signaling in the FM. Indeed, we observed an average of approximately 6-fold lower expression of the *pTCS::GFP* reporter in FMs compared to IMs (Fig. 4B, S4Ci), despite both meristems being around the same diameter under low nutrient conditions (Fig. 1E, Fig. S1D). The much lower CK signaling levels in the FM could result in changes in CK signaling having less impact on the *WUS* domain size, and could explain why reducing CK signaling further in mutant backgrounds has less effect on meristem diameter and organ number. One way to begin to test this hypothesis would be to use an inducible floral-meristem specific promoter to drive cytokinin biosynthesis to investigate whether induction of high levels of cytokinin in the FM are sufficient to cause an increased WUS domain size and floral organ number.

The analysis of organ production in *clv3* suggests the importance of the negative feedback of *CLV3* on *WUS* expression for the invariance of floral organ production within and between conditions. This property of the *CLV* feedback to add robustness to morphogenesis is a key feature of negative autoregulation motifs, which are very common motifs in gene regulatory networks and have been proposed to reduce intrinsic noise in gene expression (Alon, 2007). Variability within conditions and plasticity between conditions, notably in gene expression, are not necessarily connected (Abley *et al*., 2016). However, it appears that the *CLV* feedback in the floral meristem not only restricts variation in the number of floral organs in a given condition, it also prevents a plastic response when the growth conditions are altered (Fig. 2C, Supplementary Fig. S2B), thus showing that variability and plasticity are connected in our system. Such a correlation can also be observed in the WT inflorescence meristem where flower production rate is more variable within a condition compared to floral organ production and highly plastic in response to changes in the growth conditions.

Additional mechanisms may also be involved to ensure the invariance of floral organ production. A model has been developed to understand the preferential occurrence of tetrameric and pentameric whorls in flowers (Kitazawa and Fujimoto, 2015). One feature of this model is that later initiated primordia have a reduced strength of inhibition, which is important for regulating the number of organs per whorl. It is possible that under some circumstances, there could be a compensatory change in the size of the inhibition zone as the meristem diameter changes, which would allow a constant number of organs to be maintained. In the system characterised here, in the combined Col-0 and *clv3* data (Fig. 3C), the increase in sepal number is linear with the increase in floral meristem diameter, which suggests that in the Arabidopsis floral meristem there is no buffering mechanism that would change the size of the inhibition zone as the meristem size increases to maintain a constant number of organs. This is likely also true for the Arabidopsis IM, where it was previously shown that there is a linear correlation between SAM size and the rate of organ production for the IM for a large range of IM sizes, which would not be the case if the size of the inhibition field relative to the meristem was altering (Landrein *et al*., 2015). This does not rule out that such a buffering system exists in other species and contexts. In an evolutionary context, there may be situations when changes in meristem size are compensated for by changes in organ or inhibition zone size such that a constant floral structure can be maintained.

In the model of (Kitazawa and Fujimoto, 2015) it is also proposed that a mutual repulsion of the growth of the floral organs after initiation is necessary for the robustness of organ production in whorls. This repression could be mediated by *CUP-SHAPED COTYLEDONS* (*CUC*) genes, a family of transcription factors involved in growth control in organ boundaries (Aida *et al*., 1997; Vroemen *et al*., 2003; Nikovics *et al*., 2006; Maugarny-Calès *et al*., 2019). Mutants for *miR164c*, a microRNA involved in the silencing of *CUC1* and *CUC2* expression, show increased number of petals within flower buds (Baker *et al*., 2005) while the expression of a *miR164*-resistant *CUC2* gene also leads to the production of flowers organized following whorled-like structures on the stem (Peaucelle *et al*., 2007). The b-zip transcription factor *PERIANTHIA (PAN)*, which is expressed ubiquitously in the flower bud and promoted by *WUS* (Maier *et al*., 2009) may also be involved in this process as its mutant produces pentameric flowers in Arabidopsis without affecting the size of the floral meristems (Chuang *et al*., 1999).

Although the number of floral organs is usually invariant in the core eudicots (Endress, 2011; Ma *et al*., 2017), the flowers of *Cardamine hirsuta*, a close relative of Arabidopsis, can produce a variable number of petals. The genetic basis of this phenotype has been assessed and quantitative trait loci (QTL) controlling both mean and variability in petal number have been isolated (Pieper *et al*., 2016; Monniaux *et al*., 2018). The effect of these QTL can be suppressed by introgressing the floral homeotic gene *APETALA1* (*AP1*) of Arabidopsis into *Cardamine* thus suggesting that *AP1* can canalize floral organ production in different species (Monniaux *et al*., 2018). Although the number of petals in *Cardamine* is variable within a condition, it can also be influenced by environmental signals such as reducing the temperature, which can increase floral meristem size by delaying flower bud outgrowth and thus allow more petals to be produced (McKim *et al*., 2017). In this case, the distance available for petals to be initiated between the sepals was a key parameter controlling the number of petals produced by the flower, showing again the importance of boundary growth in the control of floral organ number. In the future, it will be important to comprehensively examine the mechanisms that modulate the level of environmental plasticity in floral organ number, beyond what we have found here for the CK signalling and *CLAVATA/WUSCHEL* circuit.

Somewhat analogous to our findings here, recent work showed that cauline and rosette axillary buds have different levels of developmental plasticity (Fichtner *et al*., 2022). Almost all cauline buds were found to activate and produce branches in a range of Arabidopsis genotypes and ecotypes, under a range of conditions, thus exhibiting invariance in their behaviour. On the other hand, it was rare for all rosette buds in a plant to activate and there was plasticity in the number of rosette branches that activated in response to photoperiod, light intensity and temperature. This suggests that the differences in plasticity between the IM and FM that we observed are just one example of how developmentally similar structures can exhibit differences in their levels of plasticity.

## Acknowledgements

We thank Hugo Tavares for help with analysis and Ottoline Leyser, Bruno Müller, Hitoshi Sakakibara and the RIKEN Institute for sharing lines. This work is supported by the Gatsby Charitable Foundation [fellowship GAT3395/DAA (to E.M.M.), AT3272/GLC (to J.C.W.L.), and GAT3395-PR4B (to H.J.)]. E.M.M. also acknowledges support from the Howard Hughes Medical Institute. Research in the laboratory of J.C.W.L. was made possible by the award of a European Research Council under the European Union’s Seventh Framework Programme (FP/2007-2013)/ERC Grant Agreement 338060. P.F.-J. acknowledges a postdoctoral fellowship provided by the Herschel Smith Foundation.

## Materials and methods

### Data

The measurements of meristem size, plastochron ratio, flower number and gene expression in IM and FM shown in Figs. 1, 3, 4 and 5 were performed on a set of data generated and partly analysed in (Landrein *et al*., 2018) (for the IM but not for the FM).

### Plant material and growth conditions

All mutant and marker lines were in the Col-0 background, which was provided by the Salk stock centre. The *pWUS*::*GFP* and *pTCSn::GFP* markers have been already described (Jönsson *et al*., 2005; Zürcher *et al*., 2013). The *ckx3-1 ckx5-2* (referred as *ckx3.5*)*, ipt3-1 ipt5-2 ipt 7-1* (referred as *ipt3.5.7*), *log4-3 log7-1* (referred as *log4.7), log1-2 log3-2 log4-3 log7-1* (referred *as log1.3.4.7*), *clv3-7* and *clv3-9* mutants have also been characterised (Fletcher *et al*., 1999; Miyawaki *et al*., 2006; Tokunaga *et al*., 2012; Nimchuk *et al*., 2015); (Melnyk *et al*., 2015).

After sterilization in a solution of 70% Ethanol + 0.01% Triton X100, all seeds were sown on plates of 0.5x MS medium (Murashige and Skoog, Sigma), placed for 3 days in a cold room for stratification and for 7 to 10 days in a short day growth chamber (8h light). The seedlings were transferred into separate pots containing soil only (Levington F2) or a mix of soil and sand (Royal horticultural society: silver sand) equalized by volume, put into the growth chamber at constant light (22°C, 160 µmol m^-2^ s^-1^) and watered with demineralized water.

### Meristem dissection and imaging

IMs were measured in plants that were a few days after bolting, when the first flowers were opening. At this stage, the main inflorescence meristem was cut a few centimeters from the tip and prepared for imaging. The cut IMs were dissected under a stereomicroscope in a box containing 1% agarose (Sigma) to remove all of the flowers older than stage 3 (Smyth *et al*., 1990) and transferred to a box of apex culture medium (ACM) supplemented with vitamins and 500 nM of BAP (6-Benzylaminopurine, Sigma) (Fernandez *et al*., 2010). To dye the cell membranes, the meristems were flooded in a solution of 0.1 mg/mL of FM4-64 (Thermo-Fisher) for 10 minutes and washed in sterile water. They were then imaged in sterile water using either a 10X or a 20X long-distance water-dipping objective mounted on an LSM700 or an LSM780 confocal microscope (Zeiss, Germany) and z-stacks of 2 μm spacing were taken. For the time-lapse microscopy of Col-0 and *clv3* meristems, the meristems were kept in ACM medium and put in a growth chamber under constant light in a closed box to prevent dehydration between imaging sessions.

### Image analysis and morphological measurements

All of the morphological analyses were performed manually using the ImageJ software (https://fiji.sc/). The size of the IM was assessed by measuring manually the distance between the meristem center and the three youngest observable initia while the plastochron ratio was assessed by measuring the mean ratio in area (measured manually on z-projections of the confocal stacks) between successive primordia ordered following the spiral pattern of phyllotaxis (Landrein *et al*., 2015). The size of the FM at stage 3, defined as a stage where sepals are visible but not yet covering the floral bud (Smyth *et al*., 1990), was measured manually, just after the emergence of the four sepals, by performing orthogonal sections along the medial and lateral axis of the flower buds (Supplementary Fig. S1A). Note that the size of the IM, measured as a radius, was converted into a diameter to ensure a correct comparison with the size of the FM, which was measured as a diameter.

The analysis of the *pTCSn::GFP* and the *pWUS::GFP* reporters in the IM was performed using a previously developed automatic pipeline (Landrein *et al*., 2018; Formosa-Jordan and Landrein, 2023) in Matlab (Mathworks Inc., Natick, MA), which is as follows. As a start, the IM is automatically detected in the maximal intensity z-projection of the fluorescence channels. To do that, an Otsu thresholding finds the IM and FMs fluorescence domains, and then, the code assumes that the IM domain is the one closest to the geometric center of the ensemble of the domains’ centroids. Then, in the sum z-projection, a fluorescence intensity profile of the IM fluorescence domain is computed as a function of the distance to the IM fluorescence domain center up to a given distance *ρ*. A generalized Gaussian is fitted to the fluorescence profile, and this allows us to extract domain features such as domain size and total fluorescence. The autofluorescence is estimated from the fit and removed in the computation of the total fluorescence. The domain size (i.e., the characteristic diameter of the expression domain) is defined as twice the distance from the domain center to the radial point where the fluorescence drops a factor *e*.

The analysis of the two reporters in FMs was performed by extending this pipeline as follows (Formosa-Jordan and Landrein, 2023). First, the contours of the stage-3 FMs are manually outlined in a maximal z-projection of the channel showing the cell membranes. Second, the FMs fluorescence domains are automatically detected in the maximal intensity z-projection of the fluorescence channels as explained above, where now the IM’s fluorescence domain is discarded. A similar analysis of the fluorescence domains is performed on the FMs, with the difference that the fluorescence profile of each FM fluorescence domain is computed up to a distance that is proportional to the size of the corresponding fluorescence domain found with the Otsu thresholding. Third, there is an alignment between the FMs outlines and the FMs fluorescence domains, such that area and fluorescence domain features for FMs are paired. This alignment is based on minimizing the distance between the centroids of the manually outlined domains and the centroids of the fluorescence domains.

### Measurements of flower and organ number

Flower production rate was assessed by measuring the number of siliques and opened flowers produced by the main inflorescence meristem within 10 days after the opening of the first flower. The production of floral organs was assessed by dissecting each day the newly opened flowers and measuring the number of floral organs they produced. This analysis was performed on 100 flowers (10 per plant) for each genotype and each independent experiment.

### Statistical analysis

All experiments except that shown in Fig. 5C were independently carried out twice and the number of biological replicates is displayed in each figure and or in its caption. The experiment shown in Fig. 5C was only performed once. Independent experiments were pooled for the measurements of inflorescence and floral meristem diameter, floral organ number, flower number, and size of the *pTCSn::GFP and pWUS::GFP* expression domain. Independent experiments were not pooled for the measurements of signals from confocal stacks of *pTCSn::GFP and pWUS::GFP* obtained by confocal microscopy as the intensity of the GFP signal can vary from experiment to experiment. All graphs and statistical analysis were performed using the R software (https://www.R-project.org). The measurements of the IM diameter, gene expression in the IM and flower production within 10 days were compared using a standard bilateral Student’s test. The measurements of FM diameter and gene expression in FMs were compared using linear mixed-effects models to take into account the fact that multiple floral meristems and flowers could come from the same plant. The measurements of floral organ number were compared by fitting a Poisson regression model and doing pairwise comparisons between conditions for each genotype, while the measurements of FM diameter and gene expression in the FM were compared by fitting a Gaussian distribution. The correlations between plastochron ratio and inflorescence meristem diameter, and between floral meristem diameter and organ number were calculated using linear models. The correlations between IM and FM diameter, and between shoot weight and IM or FM diameter were calculated using Spearman’s correlation. The plasticities of meristem diameters in response to nutrients were calculated as the percentage differences in mean meristem diameters between conditions.

## Author Contribution

J.L., H.J. and E.M.M obtained funding and supervised the work. B.L. led the project and performed the experiments. B.L. and K.A. performed the data analysis. P.F.-J. developed the tool to analyse gene expression based on confocal pictures. B.L., K.A. and J.L. wrote the paper with inputs from all co-authors.

## Data availability

All confocal images can be accessed through the Cambridge University Open Data Repository. One previously published data set (doi: 10.17863/CAM.18621) and a new data set (doi: TBC) were used. The source code and scripts for the automatic image analysis of gene expression and other analyses can be found at the Sainsbury Laboratory software repository (https: TBC).

## The following supplementary data are available

**Fig. S1.**
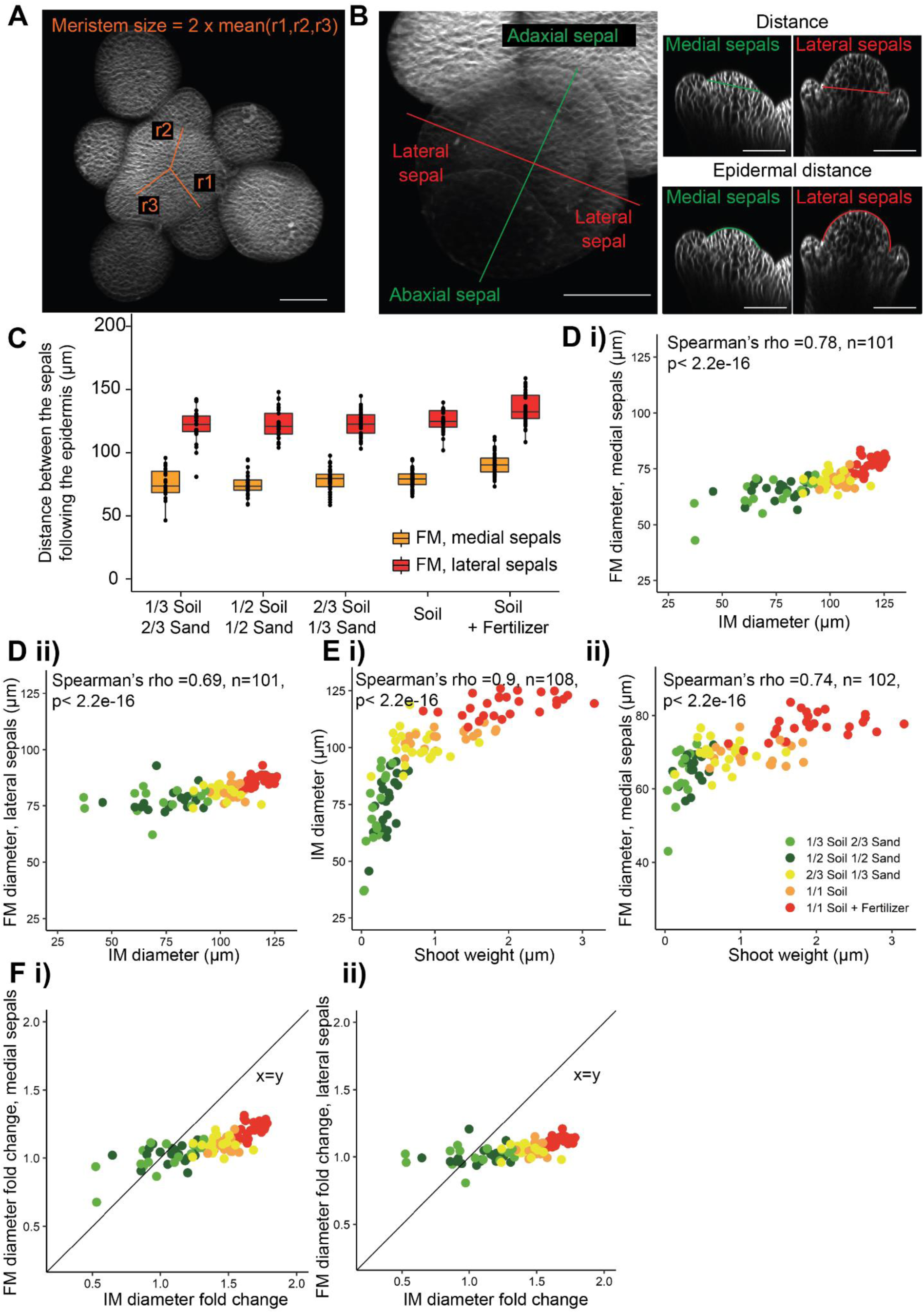
Measurements of floral meristem size in WT plants. **A.** IM size is assessed by measuring the average distance between the center of the IM and the boundary of the three youngest visible primordia. It is then converted into a diameter to allow comparison with FM size. Scale bar, 50µm. **B.** FM size is assessed by measuring the distance between medial and lateral sepals following the epidermis or in a straight line to provide two different measures, based on sections through the middle of stage 3 flowers. Scale bars, 20 µm. **C.** Distance between the medial sepals (top) and lateral sepals (bottom) following the epidermis in stage 3 flowers produced by plants growing on soils of different nutritive qualities. 23 to 43 FMs produced by 17-26 IMs from 2 independent experiments were used for each condition. **D.** Stage 3 FM diameter *vs*. corresponding IM diameter, with FM diameter assessed by measuring the distance between the medial sepals (i) and lateral sepals (ii). **E.** IM diameter *vs.* total shoot weight (i) and stage 3 FM diameter assessed as the distance between the medial sepals *vs.* total shoot weight (ii). In D and E, the plots show the combined data points from the 2 independent experiments. Where more than one FM was measured from each IM, the mean of the FM measurements is plotted. **F)** Fold change in stage 3 FM diameter relative to the average diameter of FMs in the lowest nutrient condition *vs*. corresponding fold change in IM diameter, with FM diameter assessed by measuring the distance between the medial sepals (i) and lateral sepals (ii). Colour indicates the nutrient condition, as indicated in the legend in E ii). The data on IM size (D) and shoot weight (E) were previously generated and analysed in (Landrein *et al*., 2018). The data on FM size in B, D and E were previously generated, but not analysed, in (Landrein *et al*., 2018).

**Fig. S2.**
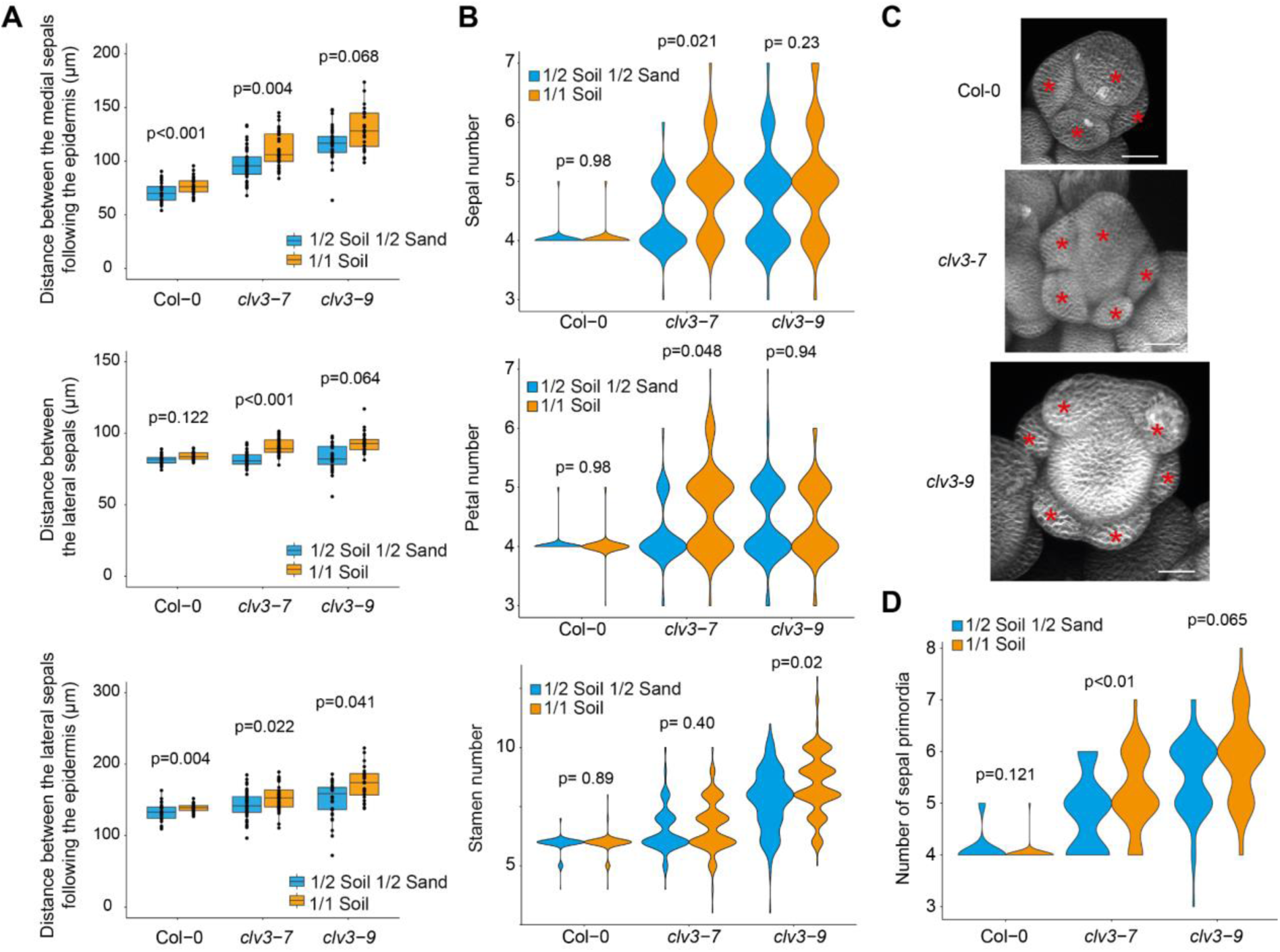
Nutrition affects floral meristem size and floral organ number in *clv3* mutants. **A.** Epidermal distance between the medial sepals, distance between the lateral sepals and epidermal distance between the lateral sepals of stage 3 flowers produced by Col-0, *clv3-7* and *clv3-9* plants. 11 to 24 plants from 2 independent experiments were sampled for each condition. A linear mixed-effect model was used to assess the effect of the growth conditions. **B.** Violin plot representation of the frequency of sepals, petals and stamens produced by Col-0 and *clv3* mutant flowers growing in different soil conditions. 200 flowers from 20 plants from 2 independent experiments were sampled for each condition. A Poisson regression model was fitted to the data and pairwise comparisons between conditions for each genotype were made. **C.** Sepal identification from confocal images of flowers close to stage 4. **D.** Violin plot representation of the number of sepal primordia in Col-0 and *clv3* mutants growing in different soil conditions. A linear mixed-effect model was used to assess the effect of the growth conditions.

**Fig. S3.**
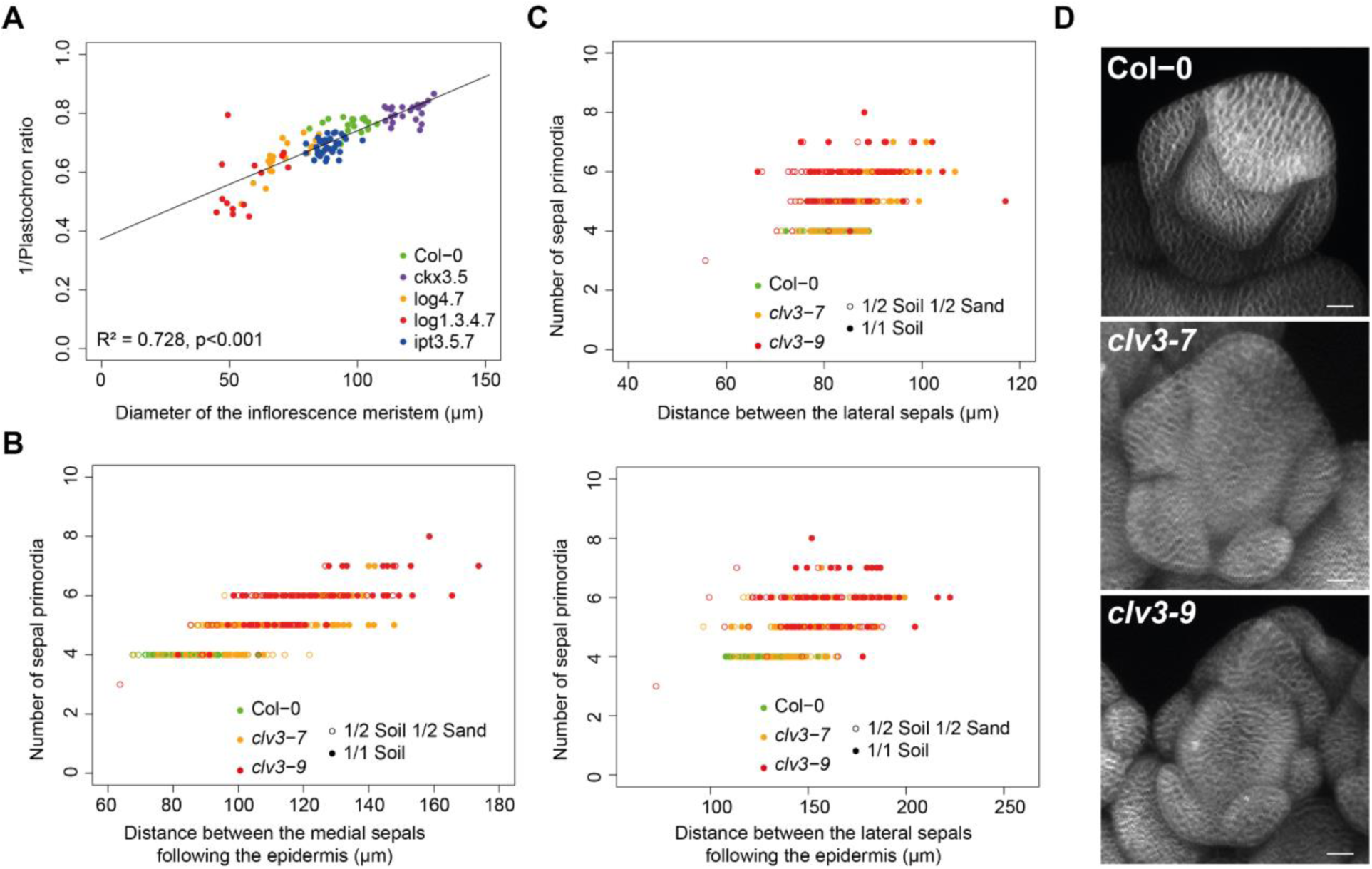
Organ number correlates to meristem size in both inflorescence and floral meristems. **A.** Correlation between the organogenesis rate (measured as the inverse of the plastochron ratio) and the size of the inflorescence meristem in Col-0 WT and in a set of mutants affected in CK metabolism. 14 to 26 IMs from two independent experiments were used for each condition. A linear model was applied to fit the data. **B.** Correlation between the number of sepal primordia and the size of the floral meristem (measured through the distance between the medial sepals following the epidermis) in a pool of Col-0 and *clv3* mutant plants growing on a mix of soil and sand or on sand only and cultivated *in vitro* for 24h. 11 to 24 plants from 2 independent experiments were sampled for each condition. **C.** Correlation between the number of sepal primordia and the size of the floral meristem (measured through the shortest distance between the lateral sepals or following along the epidermis) in a pool of Col-0 and *clv3* mutant plants growing on a mix of soil and sand or on sand only and cultivated *in vitro* for 24h. 11 to 24 plants from 2 independent experiments were sampled for each condition **D.** Representative late stage 3 primordia showing fasciation of the floral buds of *clv3-7* and *clv3-9* along the medial axis. Scale bars, 20 µm. The data on IM size and plastochron ratio (A) were previously generated and analysed in (Landrein *et al*., 2018).

**Fig. S4.**
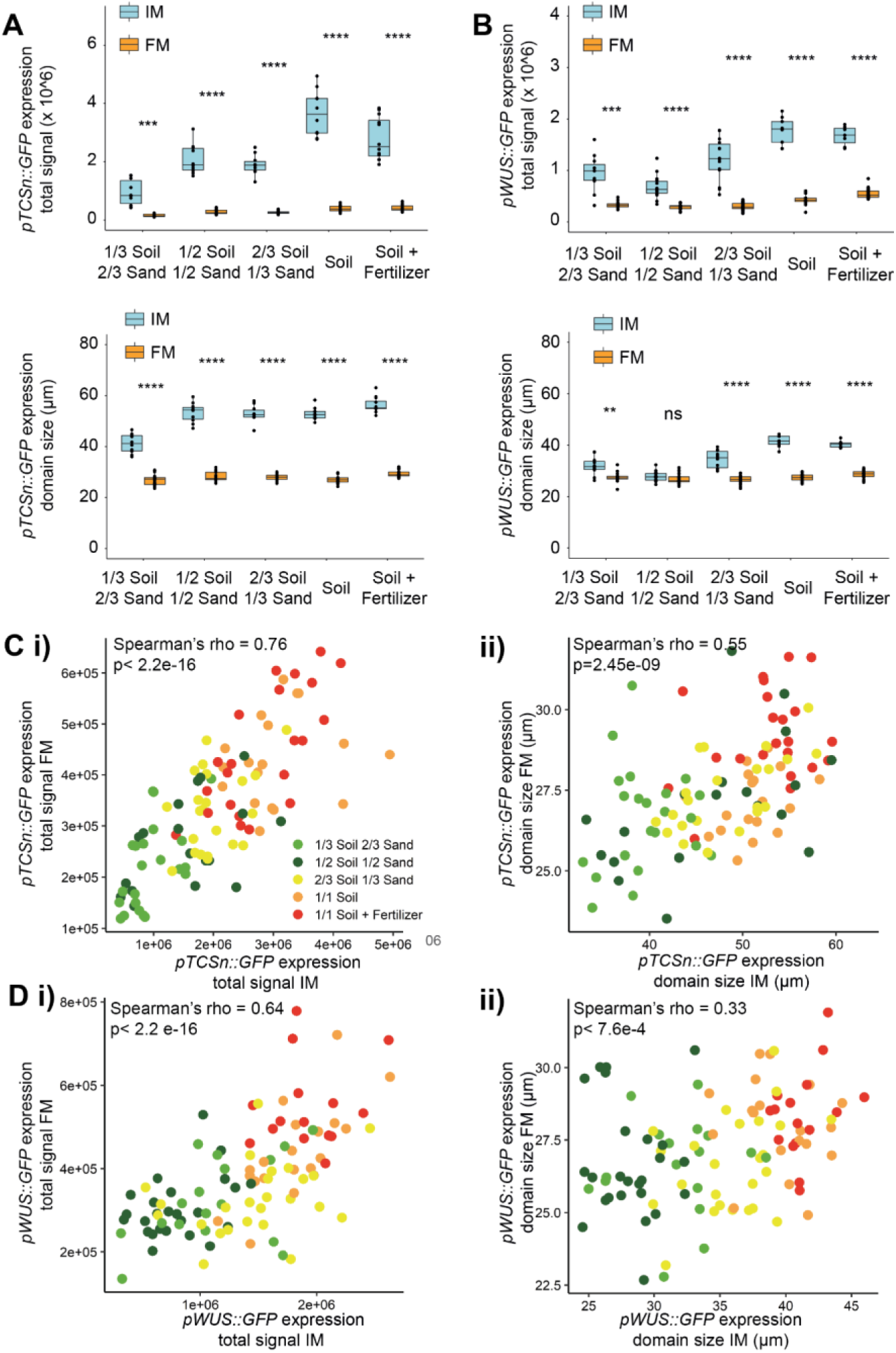
Nutrients differently affect CK response and *WUS* expression in IM and in FM of WT plants. **A.** Total signal (top) and characteristic diameter of the expression domain (bottom) of the *pTCSn::GFP* marker in stage-3 FM and in IM obtained from Col-0 WT plants growing on soils of different nutritive qualities. 12 to 22 FMs produced by 10 IMs from one experiment independent from the one shown in Fig. 4B were used for each condition. **B.** Total signal (top) and characteristic diameter of the expression domain (bottom) of the *pWUS::GFP* marker in stage-3 FM and in IM obtained from Col-0 WT plants growing on soils of different nutritive qualities. 16 to 24 FMs produced by 7 to 14 IMs from one experiment independent from the one shown in Fig. 4D were used for each condition. Meristem types were compared using bilateral Student tests. **C) i)** *pTCSn::GFP* marker total signal in stage-3 FM *vs.* corresponding IM obtained from Col-0 WT plants growing on soils of different nutritive qualities. **ii)** *pTCSn::GFP* domain diameter in stage-3 FM *vs*. IM diameter. **D) i)** *pWUS::GFP* marker total signal in stage-3 FM *vs.* IM obtained from Col-0 WT plants growing on soils of different nutritive qualities. **ii)** *WUS::GFP* domain diameter in stage-3 FM *vs*. IM diameter. In C and D, plots show the combined data points from the 2 independent experiments. Where more than one FM was measured from a given IM, the mean of FM measurements was plotted. Colour indicates the nutrient condition, as shown by legend in Ci). In A-D, total fluorescence signal measurements are in arbitrary units. The data in A-D were all previously generated and analysed for the IM but not for the FM, in (Landrein *et al*., 2018).

**Fig. S5.**
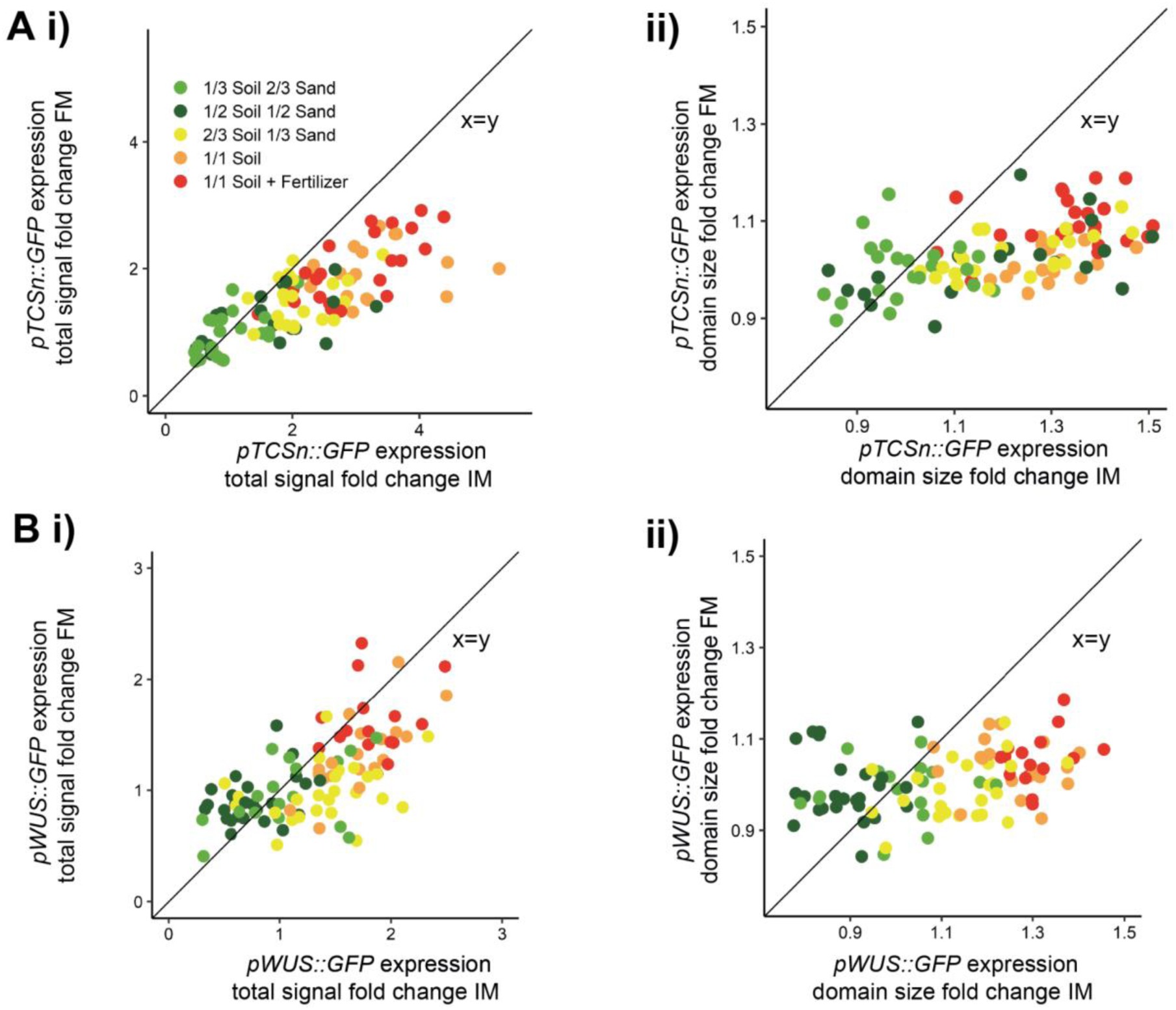
Nutrients differently affect the relative CK response and change in *WUS* expression in IM and in FM of WT plants. **A. i)** Fold change of *pTCSn::GFP* marker total signal in stage-3 FM relative to the mean *pTCSn::GFP* marker total signal in the lowest nutrient condition *vs.* corresponding fold change in IM total signal obtained from Col-0 WT plants growing on soils of different nutritive qualities. **ii)** Fold change of *pTCSn::GFP* domain diameter in stage-3 FM relative to the mean *pTCSn::GFP* domain diameter in the lowest nutrient condition *vs*. corresponding fold change in IM domain diameter. **B. i)** Fold change of *pWUS::GFP* marker total signal in stage-3 FM relative to the mean *pWUS::GFP* marker total signal in the lowest nutrient condition *vs.* corresponding fold change in IM total signal obtained from Col-0 WT plants growing on soils of different nutritive qualities. **ii)** Fold change of *WUS::GFP* domain diameter in stage-3 FM relative to the mean *WUS::GFP* domain diameter in the lowest nutrient condition *vs*. corresponding fold change in IM domain diameter. Plots show the combined data points from 2 independent experiments. Where more than one FM was measured from a given IM, the mean of FM measurements was plotted. Colour indicates the nutrient condition, as shown by legend in Ai). Total fluorescence signal measurements are in arbitrary units. The data were all previously generated and analysed for the IM but not for the FM, in (Landrein *et al*., 2018).

**Fig. S6.**
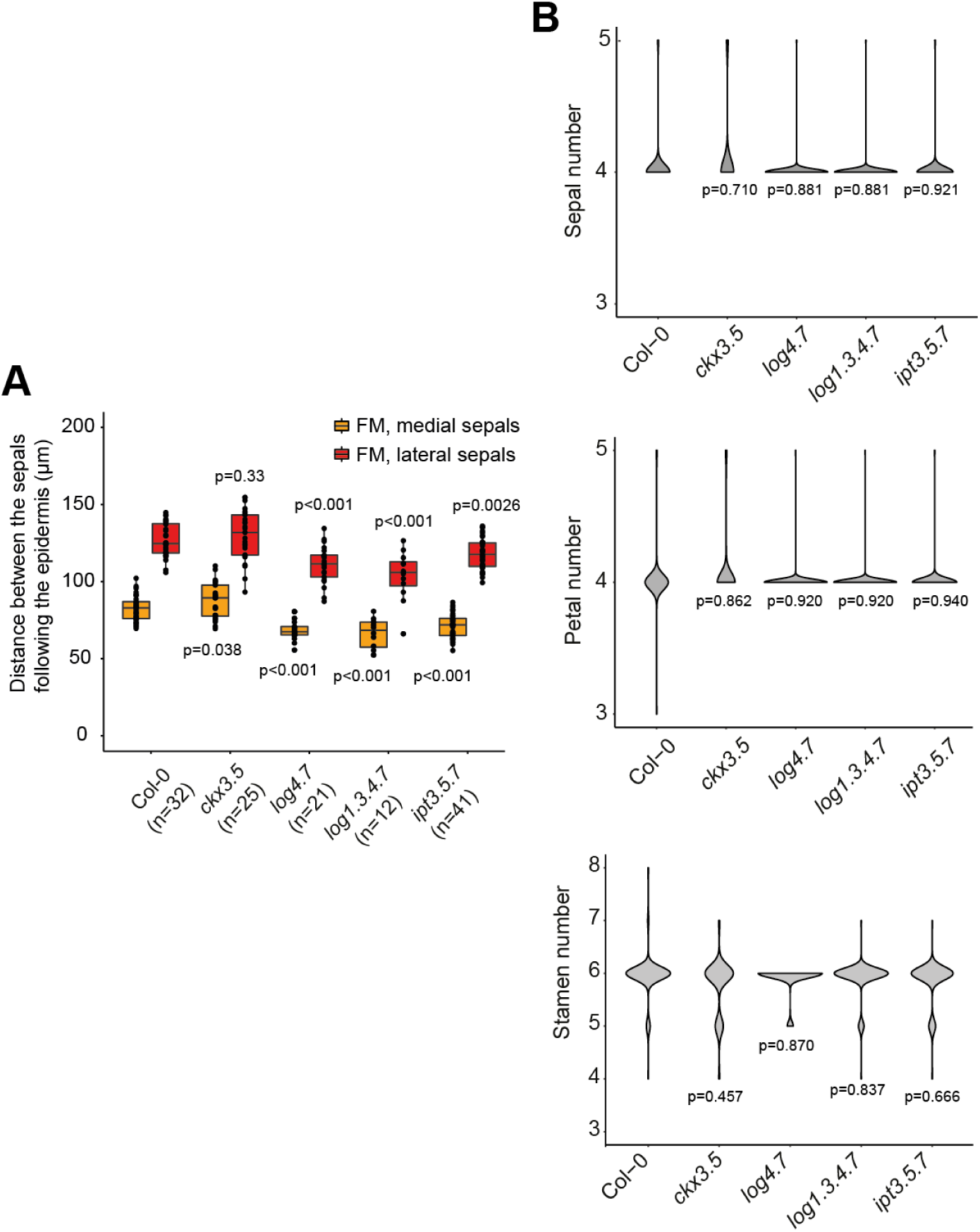
Altering CK metabolism affects inflorescence and floral meristems differently. **A.** Distance between the medial sepals and the lateral sepals following the epidermis in stage 3 flowers produced by Col-0 WT and cytokinin-associated mutant plants growing on soil. 12 to 41 FMs from 2 independent experiments were used for each condition. Each mutant was compared to Col-0 using a linear mixed-effect model. **B.** Violin plot representation of the number of sepals, petals and stamens produced by Col-0 WT and mutant plants altered in CK metabolism and growing on soil. 200 flowers from 20 plants from 2 independent experiments were sampled for each condition. Each mutant was compared to Col-0 using a linear mixed-effect model.

